# Neuroprotection in early stages of Alzheimer’s Disease is promoted by Transthyretin angiogenic properties

**DOI:** 10.1101/2021.04.16.440131

**Authors:** Tiago Gião, Joana Saavedra, José Ricardo Vieira, Marta Teixeira Pinto, Gemma Arsequell, Isabel Cardoso

**Affiliations:** i3S - Instituto de Investigação e Inovação em Saúde, Universidade do Porto, Rua Alfredo Allen 208, 4200-135, Porto, Portugal (T. Gião,; J. Saavedra,; J.R. Vieira,; M.T. Pinto,); IBMC – Instituto de Biologia Molecular e Celular, Universidade do Porto, Rua Alfredo Allen 208, 4200-135, Porto, Portugal; Faculdade de Medicina, Universidade do Porto, Alameda Prof. Hernâni Monteiro, 4200-319 Porto, Portugal; IPATIMUP – Instituto de Patologia e Imunologia Molecular, Universidade do Porto, Rua Júlio Amaral de Carvalho,45- 4200-135 Porto, Portugal; Institut de Química Avançada de Catalunya (I.Q.A.C.-C.S.I.C.), 08034 Barcelona, Spain (G. Arsequell,); Instituto de Ciências Biomédicas Abel Salazar (ICBAS), 4050-013 Porto, Portugal

**Author notes:** These authors contributed equally to this work (Tiago Gião, Joana Saavedra).

**Keywords:** Transthyretin, Alzheimer’s Disease, Basement Membrane, Angiogenesis, Neuroprotection, Chick Chorioallantoic Membrane Assay, TTR tetramer stabilizers

## Abstract

While still controversial, it has been demonstrated that vascular defects can precede the onset of the other AD hallmarks features, making it an important therapeutic target. Given that the protein transthyretin (TTR) has been established as neuroprotective in AD, here we investigated the influence of TTR in the vasculature. AD transgenic mice with TTR genetic reduction, AD/TTR+/−, exhibited a thicker BM in brain microvessels and decreased vessel length than animals with normal TTR levels, AD/TTR+/+. Further *in vivo* investigation, using the chick chorioallantoic membrane (CAM) assay, revealed that TTR is a pro-angiogenic molecule. Also, TTR increased the expression of key angiogenic molecules, by endothelial cells under tube formation conditions. We showed that TTR reduction leads to a thicker BM in AD mice than in NT animals, strengthening the idea that TTR is a neuroprotective protein. We also studied the effect of TTR tetrameric stabilization on BM thickness, showing that AD mice treated with iododiflunisal (IDIF) displayed a significant reduction of BM thickness and increased vessel length when compared to non-treated littermates. Our *in vivo* results show the involvement of TTR in angiogenesis, particularly as a modulator of vascular alterations occurring in AD. Since TTR is decreased early in AD, its tetrameric stabilization can represent a therapeutic avenue for the early treatment of AD through the maintenance of the vascular structure.

## Introduction

Alzheimer’s Disease (AD) patients undergo several neurovascular changes at different levels. Brain vascular dysregulation is the earliest and strongest factor during the disease progression and is followed by amyloid-β (Aβ) peptide deposition, glucose metabolism dysregulation, functional impairment and gray matter atrophy, in this order [1]. Decreased expression of the low-density lipoprotein receptor-related protein 1 (LRP-1) and P-glycoprotein (P-gp), as well as up-regulation of the receptor for advanced glycation end products (RAGE), are mechanisms reported to be changed in AD patients, leading to Aβ accumulation in the brain [2,3]. In addition to defective clearance mechanisms, increased endothelial pinocytosis, decreased number of mitochondria, decreased glucose transporter (GLUT)-1 and loss of tight/adherents junctions are features detected in AD [4]. The reduction of the capillary density is also characteristic of the AD brains [5]. This is due to an aberrant angiogenesis with premature pruning of capillary networks. This defective angiogenesis may be caused by a lack of angiogenic stimuli and unresponsive endothelium [6]. Although other authors describe increased vascular density in AD [7], the underlying angiogenic process has pathological characteristics. Some studies suggest that promotion of angiogenesis results in concomitant BBB disruption and vessel leakiness [7]. Other studies defend that the vascular damage is a consequence of poor blood perfusion of the brain, leading to hypoperfusion/hypoxia causing the BBB dysfunction [8]. Other authors argue that the accumulation of Aβ in the walls of the capillaries can contribute to the reduced brain capillary density in AD via anti-angiogenic activity [9,10]. Another observed alteration in AD is the increased thickness of the vascular BM in AD [11]. Since the increase in BM thickness occurs before Aβ deposition, it is speculated that it functions as a physical barrier to the Aβ clearance across the BBB [12]. Some studies have related this BM thickening with increased collagen IV content, in AD and ageing [13,14].

Transthyretin (TTR), a 55 kDa homotetrameric plasma and cerebrospinal fluid (CSF) protein, transports retinol through binding to the retinol-binding protein (RBP), which binds at the surface of TTR, and thyroxine (T4), which binds at a central hydrophobic channel formed at the dimer-dimer interface [15]. In the CSF, TTR is the main Aβ binding protein [16], providing neuroprotection by avoiding Aβ aggregation [17–24] and toxicity [17,25], and by participating in Aβ brain efflux at the BBB [26]. TTR is early decreased in AD, both in plasma [27–29] and in the CSF [30], probably due to its tetrameric instability [27,31], hypothesized to result in accelerated clearance and lower levels. TTR instability is also a key feature in familial amyloid polyneuropathy (FAP), a systemic amyloidosis that is usually caused by mutations in TTR. The amyloidogenic potential of the TTR variants is inversely correlated with its tetrameric stability [32], and the dissociation of the tetramer into monomers is at the basis of the events that culminate with TTR amyloid formation [33,34]. TTR stabilization, used as a therapy in FAP [35,36], can be achieved through the use of small-molecule compounds sharing molecular structural similarities with T4 and binding in the T4 central binding channel [37–39]. Although no TTR mutations have been found in AD patients [22], TTR stabilization has also been proposed as a therapeutic strategy to recover its ability to protect in AD [19,40], and shown beneficial in a mouse model of AD [40,41]. Iododiflunisal (IDIF), a potent TTR stabilizer, was administered to AD mice and bound plasma TTR displacing T4, resulting in decreased Aβ amyloid burden and total Aβ brain levels, and improved cognition [41]. Interestingly, TTR stabilization by IDIF improves TTR-assisted Aβ brain efflux in vitro and enhanced the expression of LRP-1 in vivo [31]. The formation of TTR-IDIF complexes enhances BBB permeability of both IDIF and TTR, in vivo [42].

TTR has also been implicated in angiogenesis and the first reports of its involvement have been described in diseases such as FAP [43]; in diabetic retinopathy [44,45], and lately, in cancer [46]. As reported, a study investigated the effect of TTR in angiogenesis by treating human umbilical vein endothelial cells (HUVECs) with wild-type (WT) TTR or a common FAP TTR mutant, V30M. The authors concluded that the TTR mutant inhibited cell migration and decreased survival relative to the WT TTR, by down-regulating several pro-angiogenic genes for angiopoietin-2 (Ang-2), VEGF receptors 1 and 2, basic fibroblast growth factor (bFGF) and transforming growth factor-beta 2 (TGF-β2) [43]. In another study, to investigate how TTR affects the development of new vessels in diabetic retinopathy (DR), human retinal microvascular endothelial cells (hRECs) were cultured with TTR in natural and simulated DR environments (hyperglycemia and hypoxia). In the DR environment, TTR inhibited cell proliferation, migration and tube formation, by repressing the expression of the pro-angiogenic genes Ang-2 and VEGF receptors 1 and 2 [44]. Conversely, in a low glucose environment, these angiogenesis-related features were improved by TTR. Recently, it was reported that TTR levels were increased in human serum of lung cancer patients. Additionally, TTR was shown able to promote tumour growth by enhancing several lung ECs functions as permeability, migration and tube formation [46]. However, TTR potential in angiogenesis has never been addressed *in vivo* and the possible participation of TTR in brain angiogenesis and vascular alterations has never been elucidated.

Taking these evidences into account, this work aimed at investigating the angiogenic potential of TTR and at assessing its involvement in the vascular impairment that occurs in AD.

## Material and Methods

### Animals

Two mouse models were used in this work, an AD transgenic and a non-transgenic (NT) mouse models, both established in different TTR genetic backgrounds.

The AD mouse model AβPPswe/PS1A246E/TTR was generated by crossing the AD mouse model AβPPswe/PS1A246E [47] (B6/C3H background) purchased from The Jackson laboratory with TTR-null mice (TTR−/−) (SV129 background) [48] as previously described [49]. F1 animals AβPPswe/TTR+/− and PS1A246E/TTR+/− were crossed to obtain AβPPswe/PS1A246E/ TTR+/+, AβPPswe/PS1A246E/TTR+/−, AβPPswe/ PS1A246E/TTR−/− and NT controls NT/TTR+/+, NT/TTR+/− and NT−/−. The colony was maintained on a B6/C3H/SV129 genetic background. Hereafter, the AβPPswe/PS1A246E/TTR colony will be referred to as AD/TTR, and the different genotypes AβPPswe/PS1A246E/TTR+/+, AβPPswe/PS1A246E/TTR+/−, and AβPPswe/PS1A 246E/TTR−/− referred to as AD/TTR+/+, AD/ TTR+/−, and AD/TTR−/−, respectively. Animals were housed in a controlled environment (12-hour light/dark cycles, temperature between 22-24°C, humidity between 45–65% and 15-20 air changes/hour), with freely available food and water. All the above experiments were approved by the Institute for Research and Innovation in Health Sciences (i3S) Animal Ethics Committee and in agreement with the animal ethics regulation from Directive 2010/63/EU.

In order to study the role of TTR in collagen IV deposition or in vessel density, cohorts of littermates 7-month-old female mice AD/TTR+/+ (n=7) and AD/TTR+/− (n=7), cohorts of littermates 3-month- old female mice NT/TTR+/+ (n=4) and NT/TTR+/− (n=4) and one cohort of 3-month-old female mice AD/TTR+/− were used. AD/TTR+/− female control mice (n=6) or treated with IDIF (n=6) [41] for two months (from 5 to 7-month-old), were used to investigate the relevance of TTR stabilization in collagen type IV levels, in AD.

### Collagen IV Immunohistochemistry

Free-floating 30 μm-thick coronal brain sections of mice were permeabilized with 0.25% Triton X-100 in phosphate-buffered saline (PBS) for 10 min at room temperature (RT), blocked with 5% bovine serum albumin (BSA) in PBS for 1 hour at RT and incubated with primary rabbit anti-collagen IV antibody (1:100) (Abcam) in 1% BSA in PBS overnight at 4°C. Next, sections were washed with PBS and incubated with Alexa Fluor-568 goat anti-rabbit IgG antibody (1:2000) for 1 hour at RT. All steps were performed with agitation. To remove tissue autofluorescence, sections were covered with Sudan black B solution (0.3% Sudan black B in 70% ethanol) applied for 5 minutes at RT, followed by multiple washing steps with PBS at RT with agitation. The brain sections were dried for 20 minutes at RT and mounted on 0.1% gelatin-coated slides with Fluoroshield^TM^ with DAPI (Sigma-Aldrich). Sections were visualized and photographed using a Zeiss Axio Imager Z1 microscope equipped with an Axiocam MR3.0 camera and Axivision 4.9.1 software. A total of twenty randomly selected vessels in the hippocampus and/or cortex of each mouse was photographed at 100x magnification, and the ratio intensity/area was measured using the ImageJ software.

To assess the vascular density of mice brains, 30 μm-thick coronal brain sections were boiled at 90 ° C in citrate buffer for 15 minutes for antigenic recovery and then washed with 0.3% Triton X-100 in PBS for 10 minutes at RT. Tissues were blocked/permeabilized with a solution of 1% BSA and 0.5% Triton X-100 in PBS, overnight at 4°C. The coronal sections were then incubated for 72 hours at 4°C with primary rabbit anti-collagen IV antibody (1:200) (Abcam) in a solution with 1% BSA, 0.5% Triton X-100 and 2% fetal bovine serum (FBS) in PBS. After, tissues were washed with 0.3% Triton X-100 in PBS at 4°C. Next, sections were incubated with Alexa Fluor-568 goat anti-rabbit IgG antibody (1:500) overnight at 4°C, followed by washing with 0.3% Triton X-100 in PBS and then dried for 20 minutes at RT and mounted on silane pre-coated slides with FluoroshieldTM with DAPI (Sigma-Aldrich). Sections were visualized and photographed using a Zeiss Axio Imager Z1 microscope equipped with an Axiocam MR3.0 camera (Carl Zeiss) and Axiovision SE64 Rel. 4.9.1 software. A total of twenty-twenty five fields of view were randomly selected from each brain section and photographed at 20x magnification. The total length of the blood vessels per field were measured using the ImageJ software.

### Production and purification of human recombinant TTR

Human recombinant WT TTR (rec TTR) was produced in a bacterial expression system using *Escherichia coli* BL21 [50] and purified as previously described [51]. Briefly, after growing the bacteria, the protein was isolated and purified by preparative gel electrophoresis after ion exchange chromatography.

### Purification of human TTR from sera

Human plasma from donors who were informed of the purpose of the study and gave their written consent, were collected in accordance with the approved guidelines. Samples were subjected to affinity chromatography to isolate human TTR (hTTR); for this we used 1 mL column of NHS-activated Sepharose coupled to rabbit anti-human TTR (Dako). The column was washed with PBS and then incubated with 500 μL of human plasma for 2 hours at RT. To elute TTR from the column, 5 mL of Gentle Ag/Aβ elution buffer (Thermo Scientific) were applied, and 1 mL-aliquots were collected and OD 280 nm was registered.

### Cell culture

The immortalized human cerebral microvascular endothelial cell line, hCMEC/D3 (Tebu-Bio) is a well-characterized *in vitro* model of BBB. The hCMEC/D3 cells were used between passage 25 and 35 and cultured following the available data sheet. All culture flasks were coated with rat tail collagen type I solution (Sigma) at a concentration of 150 μL/mL and were incubated for 2 hours at 37°C. Cells were cultured in EBM-2 medium (Lonza) containing 5% FBS (Gibco), 1% of penicillin-streptomycin (Lonza), 1.4 μM of hydrocortisone (Sigma-Aldrich), 5 μg/mL of ascorbic acid (Sigma-Aldrich), 1% of chemically defined lipid concentrate (Gibco), 10 mM of 4-(2-hydroxyethyl)-1-piperazine-1-ethanesulfonic acid (HEPES) (Gibco) and 1 ng/mL of human bFGF (Sigma-Aldrich). Cells were incubated at 37 °C in a humidified atmosphere with 5% of CO_2_. Cell culture medium was changed every 2–3 days.

### Tube formation assay

hCMEC/D3 cells, grown in 25 cm^2^ flasks, at a confluence of 80-90% were incubated for 24 hours with EBM-2 medium (Lonza) containing 1% FBS (Gibco) (negative control), with bFGF (35 ng/mL) (positive control), or with rec TTR at different concentrations (10, 25, 250, 500 nM and 1 μM) or with hTTR 250 nM. Then, cells were transferred into 96-well plates, previously coated with Matrigel (Corning), and grown in the same conditions of media, bFGF or TTR for another 9 hours. Then, cells were photographed using the In Cell Analyzer 2000 (GE Healthcare) (magnification ×10). The supernatants were collected, centrifuged at 14.000 rpm for 10 minutes and stored at −20°C. Each condition was performed in triplicate and experiments were repeated three times.

### Quantification of angiogenesis-related proteins

The angiogenesis-related proteins interleukins 6 and 8 (IL-6, IL-8), angiopoietin 1 and 2 (Ang-1, Ang-2), epidermal growth factor (EGF), basic fibroblast growth factor (bFGF), platelet endothelial cell adhesion molecule (PECAM-1), placental growth factor (PlGF), VEGF and tumor necrosis factor α (TNF-α) were quantified in the supernatants collected from hCMEC/D3 grown under conditions of tube formation in the presence of media alone or with 1 μM rec TTR, using the LEGENDplex Human Angiogenesis Panel (BioLegend) bead-based immunoassay. The assay was performed according to the manufacturer’s recommendations. Analysis was performed using a BD Accuri C6 (BD Biosciences) and LEGENDplex^TM^ Data Analysis software v8.0 (BioLegend).

### ELISA analysis for IL-6

IL-6 was also quantified in the supernatants collected from hCMEC/D3 cells used for the tube formation, in the presence of media alone or with rec TTR at different concentrations (10, 25, 250 nM and 1 μM), using a LEGEND MAX™ Human IL-6 Sandwich Enzyme-Linked Immunosorbent Assay (ELISA) Kit (BioLegend) with pre-coated plates. The assay was performed according to the manufacturer’s recommendations. Analysis was performed using Synergy Mx and by measuring absorbance at 450 and 570 nm. A standard curve was generated for IL-6 from 7.8 pg/mL to 500 pg/mL.

### Angiogenesis chick chorioallantoic membrane (CAM) assay

Commercially available fertilized chick (*Gallus gallus*) eggs were horizontally incubated at 37 °C, in a humidified atmosphere. On embryonic development day (EDD) 3, a square window was opened in the shell after removal of 1.5–2 mL of albumen, to allow detachment of the developing CAM. The window was sealed with a transparent adhesive tape and eggs re-incubated. On EDD10, rec TTR (1 μM), hTTR (1 μM), PBS (vehicle, negative control) and bFGF (50 ng/μL, positive control) were placed on top of the CAM, into 3 mm silicone rings, under sterile conditions (1 condition per egg). Eggs were re-sealed and returned to the incubator for an additional 72 hours. On EDD13, rings were removed, the CAM was excised from embryos and photographed *ex-ovo* under a stereoscope, using a 20× magnification (Olympus, SZX16 coupled with a DP71 camera). The number of new vessels (< 20 μm diameter) growing radially towards the inoculation area was counted in a blind fashion manner.

### In vivo analysis of vascular permeability

The CAM model was also used to evaluate vascular permeability or vessel leakage, as measure of TTR induced neo-vessels functionality. Embryos were cultured *ex ovo*. To prepare shell-less CAM, eggs were incubated as described above and on EDD3, the content of the egg was transferred to sterile weigh boats, covered with square petri dishes and returned to the incubator for additional 7 days. At EDD10, 10 μl of PBS, rec TTR (1 μM) and VEGF (4 ng/μL) were inoculated on distinct sites of the same egg, twice each, into 3mm silicone rings under sterile conditions. Three independent experiments were performed summing a total 16 CAM sites/condition (8 eggs). After 3 days (EDD13), chicken embryos were injected intravenously with 100 ul of Even’s Blue Dye (EBD, Sigma) solution (0.5% EBD, 5% BSA in PBS) and further incubated for 60 minutes. After incubation, embryos were perfused with saline. The tissue underlying the rings (inoculation sites) was removed, cleaned in saline, blotted dry, weight, homogenized and incubated in 200 ul of formamide (Sigma), at 38ª for 48h, to release the extravasated dye. The samples were centrifuged and 175 μl of supernatant was quantified spectrophotometrically at 620 nm. The amount of EBD in the experimental samples was calculated by interpolating to a standard curve and the concentration of EBD per g of tissue was determined.

### Statistical Analysis

All quantitative data were expressed as mean ± standard error of the mean (SEM). Initially, data was assessed whether it followed a Gaussian distribution. In the cases of non-Gaussian distribution comparisons between two groups were made by non-parametric Kruskal-Wallis test and comparisons between two groups were made by Student t-test with a Mann Whitney test.

When found to follow a Gaussian distribution, differences among conditions or groups were analyzed by one-way ANOVA with the appropriate post hoc pairwise tests for multiple comparisons tests. Differences in CAM assay and IL-6 Elisa kit were analyzed using one-way ANOVA followed by Tukey’s multiple comparison test. P-values lower than 0.05 were considered statistically significant. Statistical analyses were carried out using GraphPad Prism 8 software for Windows.

## Results

### TTR influences vascular features in the mouse brain

For this work we have used AD/TTR+/+, AD/TTR+/−, NT/TTR+/+ and NT/TTR+/− animals. We did not analyze the respective TTR−/− animals, although we obtained them in the course of breeding, to avoid indirect effects of TTR-deficiency including compensatory processes, that could confound our interpretations. Additionally, the TTR+/− animals, in particular the AD/TTR+/−, are a better representation of the behavior of TTR in AD, since TTR is decreased in this pathology but not absent.

#### Reduction of TTR increases the thickness of the collagen IV layer in brain microvessels of AD mice

To investigate a possible relation of TTR reduction with the thickening of the BM and with the structural vascular alterations reported in AD, we evaluated collagen IV levels in brain microvessels, in the AD/TTR mouse model. This model established in different TTR genetic backgrounds [49], bears two AD-related transgenes (APP and PSEN1) and Aβ deposition starts at around 6 months [52]. In comparison to males, females present a more severe form of AD-like disease, thus in this study we used 7-month-old female animals. AD/TTR+/− females were compared to littermates with normal TTR expression, AD/TTR+/+. Our results revealed a significantly thicker collagen IV layer in 7-months old AD/TTR+/− as compared to AD/TTR+/+ animals, as analysed in cortex microvessels (Figure 1.A1). Altogether, our results implicate TTR in vascular processes, which are known to be early dysregulated in AD.

**Figure 1.**
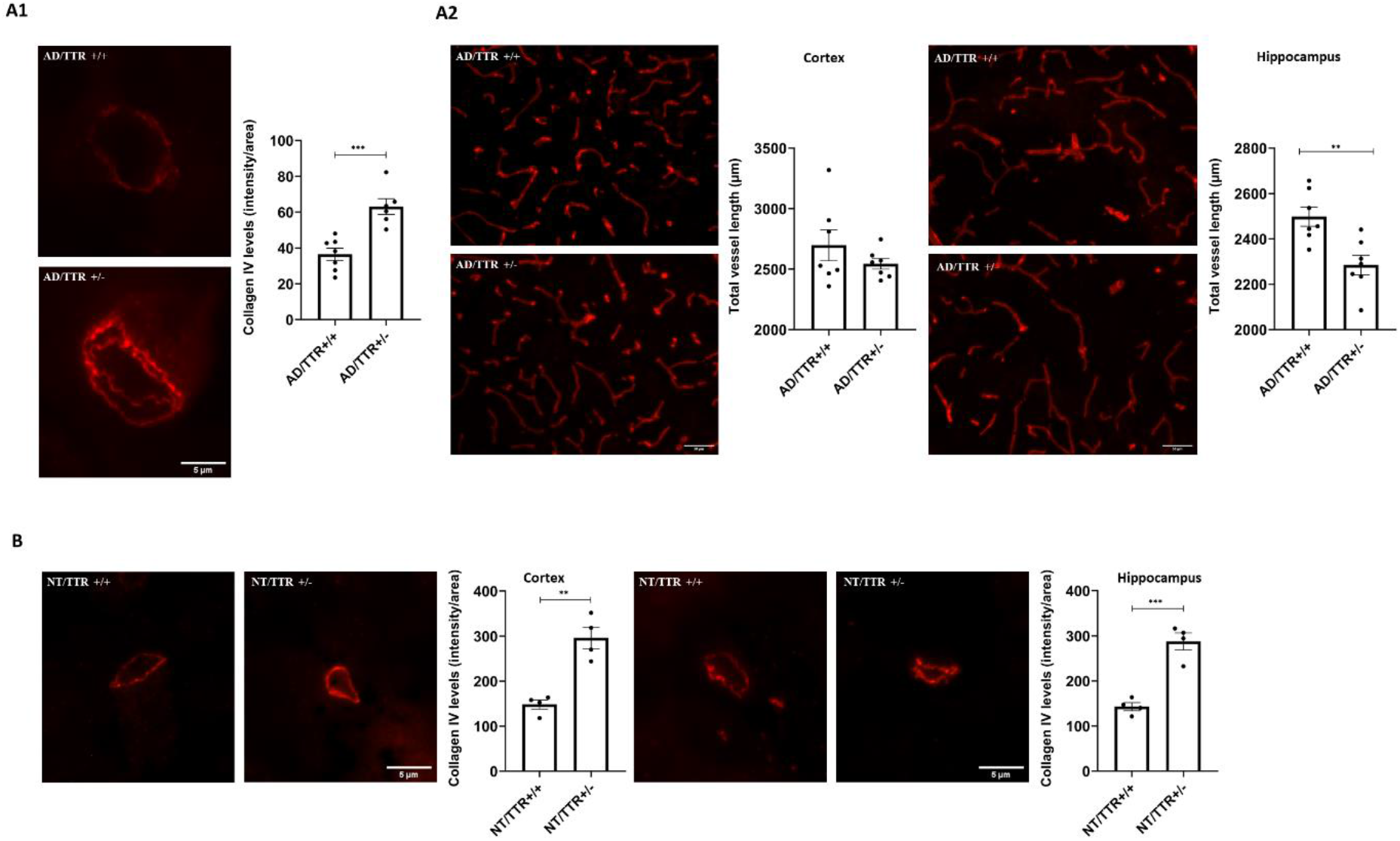
Effect of TTR reduction in vascular features of brain microvessels. **(A1)** Representative images and quantification plots of collagen IV levels in the BM of cortex vessels derived from 7-month-old AD mice with different TTR genetic backgrounds, AD/TTR/+/+ (n=7) and AD/TTR+/− (n=6), showing significantly increased levels in microvessels from AD/TTR+/− compared to AD/TTR+/+ mice. Scale bar = 5 μm. **(A2)** Representative images and quantification plots of length and area of brain vessels, as evaluated by collagen IV staining, from 7-month-old AD mice with different TTR genetic backgrounds, AD/TTR/+/+ (n=7) andAD/TTR+/− (n=7), showing significantly decreased vessel length in the hippocampus of AD/TTR+/− compared to AD TTR+/+. Scale bar = 50 μm. **(B)** Representative images of the cortex and hippocampus and quantification plots of collagen IV immunostaining in microvessels of NT/TTR+/+ and NT/TTR+/− 3-month-old mice. An increase in collagen IV content in NT/TTR+/− mice (n=4) relative to NT/TTR+/+ littermates (n=4) is observed. Scale bar = 5 μm. Data are expressed as mean ± SEM. ** p<0.01; ***p<0.001.

#### Reduction of TTR decreases the length of brain microvessels in AD mice

To understand if TTR affects other cerebrovascular features and if the effect observed at the level of the BM is related to angiogenesis, we measured brain vascular density in the same animals.

Both cortex and hippocampus were analyzed and our results show that, in hippocampus, reduction of TTR resulted in decreased vessel length in AD/TTR+/− mice as compared to AD/TTR+/+ (Figure 1.A2, right panels). In the cortex, the differences were not statistically significant (Figure 1.A2, left panels), although there was always a pattern of reduction of the length, as TTR is reduced. These observations support the results obtained for the BM thickness and further implicate TTR in angiogenesis, especially in the hippocampus, a particularly relevant brain area in the initial stages of AD.

#### Reduction of TTR increases the thickness of the collagen IV layer in brain microvessels of young non-transgenic mice

Although our results suggest that TTR influences the thickness of the BM, in particular the collagen IV layer, we could not determine if the effect was direct or indirect. One hypothesis is that high levels of Aβ, as it happens in AD, either due to increased production, reduced elimination or both, could be responsible for the increase in collagen IV. It is possible that AD/TTR+/− mice show increased amount of collagen IV because less TTR is available to interact with and to eliminate Aβ. Thus, in order to unravel this question, we compared collagen IV levels in non-transgenic (NT) littermate mice with two different TTR backgrounds, NT/TTR+/+ and NT/TTR+/− allowing to understand if TTR is directly involved. Furthermore, this evaluation was performed at the age of 3 months, which in the AD background is prior to amyloid deposition [52]. Results presented in Figure 1.B clearly show that NT/TTR+/− mice presented significantly higher levels of collagen IV in brain microvessels of both the cortex and hippocampus, as compared to NT/TTR+/+ animals, thus suggesting that it is, in fact, a direct effect of TTR.

### TTR possesses angiogenic activity

#### TTR promotes tube formation by hCMEC/D3 cells

The tube formation assay is a powerful *in vitro* test encompassing EC adhesion, migration, protease activity and tube formation (capillary-like structures). Thus, and to explore the angiogenic activity of TTR, endothelial cells of human brain origin, hCMEC/D3 cells, grown under tube formation-conditions, on Matrigel, were incubated with different concentrations of rec TTR. The results are displayed in Figure 2.A, and reveal that TTR affects the tube formation processes in a dose dependent-manner. Concentrations of TTR equal or above 250 nM result in a significantly higher area covered by the capillary-like structures, as compared to the negative control. These TTR concentrations are well below its physiologic concentration in plasma, and are similar to its concentration in the CSF. To confirm that the angiogenic effect was indeed provided by TTR, human TTR isolated from serum (hTTR) was also evaluated, corroborating the angiogenic activity of TTR (Figure 2.A’).

**Figure 2.**
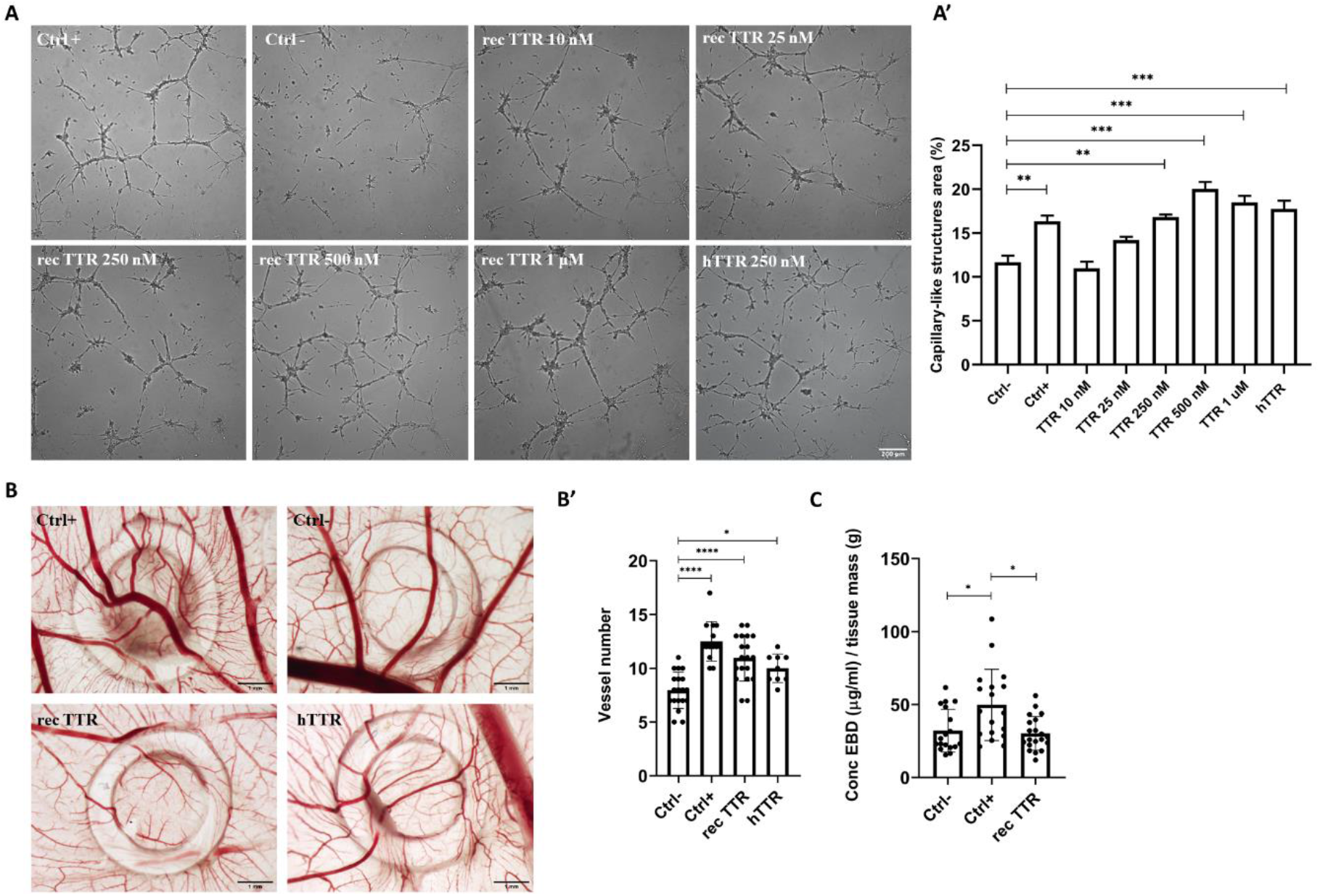
TTR angiogenic activity. **(A)** Representative images of tube formation by hCMEC/D3 cells. Cells were plated on Matrigel in the absence (negative control, Ctrl-) or presence (positive control, Ctrl+) of bFGF (35 ng/mL) or with TTR at different concentrations (10 nM-1 μM). Scale bar = 200 μm. **(A’)** The quantification plot shows that TTR concentrations equal or above 250 nM result in a significantly higher area covered by the capillary-like structures, than in the negative control. **(B)** Representative images of the chick chorioallantoic membrane (CAM) assay. **(B’)** Quantification plot of the number of new vessels (< 20 um) growing towards the inoculation site, delimited by the ring mark, induced by PBS), (Ctrl−, n=18), basic fibroblast growth factor (bFGF, 50 ng/μL) (Ctrl+, n=14), human recombinant TTR (rec TTR, 1 μM, n=19) or TTR isolated from human plasma (hTTR, 1 μM, n=9). TTR, both rec TTR and hTTR, had a significantly higher angiogenic response than the negative control. Scale bar = 1 mm. **(C)** *In vivo* vascular permeability was measured in CAM model by quantification of leaked EBD. The permeability of the new vessels induced by TTR (n=20) was similar to the negative control (PBS, n=18), in contrast to the significantly higher permeability of vessels induced by VEGF (n=18). Data are expressed as mean ± SEM. * p<0.05; ** p<0.01; ***p<0.001; ****p<0.0001.

#### TTR is angiogenic *in vivo* and the neovessels formed are functional

To further confirm the angiogenic activity of TTR, we used the CAM assay. Both rec TTR and hTTR were tested and at 1 μM induced a significantly higher angiogenic response than the negative control, as deduced by the higher number of detected new vessels (vessels with a diameter under 20 um) (Figure 2.B and 2.B’). TTR angiogenic response was comparable to the positive control (bFGF), in particular for the rec TTR.

Also using the CAM in vivo model, we studied the permeability of the TTR-induced vessels, by quantifying the leakage of EBD. This assay indicated that the permeability of TTR-induced vessels is comparable to the negative control (PBS), and significantly different from the positive control (VEGF) (Figure 2.C). It can be inferred that TTR induced neo vessels are functional (in contrast with leakier vessels induced by VEGF).

### TTR regulates angiogenic molecules

To further explore the molecular mechanisms underlying the angiogenic activity of TTR, supernatants of hCMEC/D3 cells grown under tube-formation conditions, in the presence of rec TTR (1 μM) or with media alone, were used to identify key targets involved in angiogenesis which could be affected by TTR.

Among the ten molecules analyzed, IL-6, IL-8, Ang-2 and VEGF were differentially overexpressed in the presence of TTR, whereas the remaining six presented concentrations below the detection limit. As shown in Figure 3.A, the expression of detected molecules was significantly increased relative to the negative control when stimulated with rec TTR (1μM) indicating that TTR acts as a pro-angiogenic molecule by increasing the expression of those molecules. It is possible that TTR affects other angiogenic molecules, possibly even those that were undetected by the current approach.

**Figure 3.**
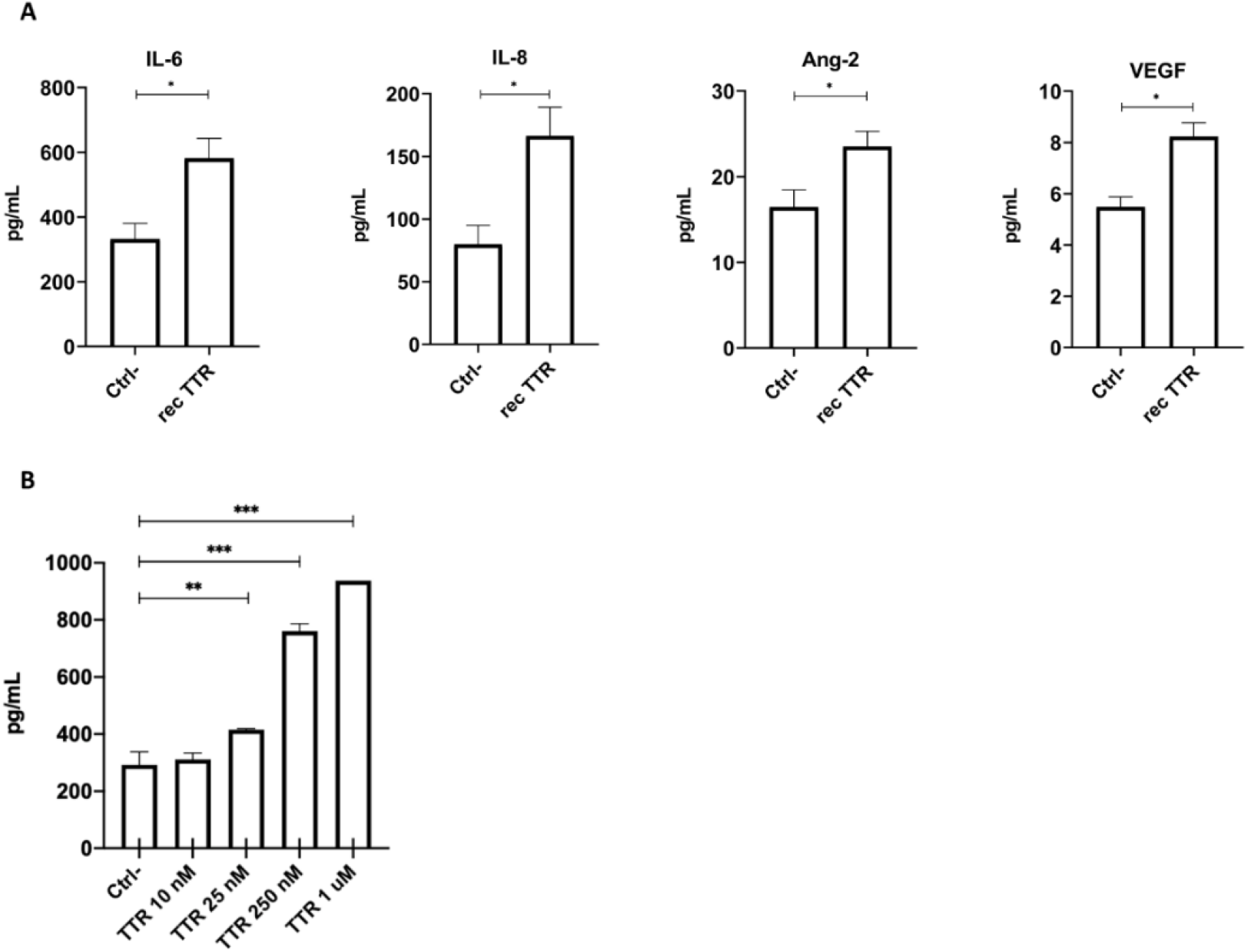
Quantification of angiogenesis-related proteins. Supernatants from hCMEC/D3 cells grown under conditions of tube formation were collected 9h after incubation with media alone or with rec TTR (1 μM). (**A)** Ten of the most common angiogenesis-related proteins were quantified using bead-based LEGENDplex assay by flow cytometry. Rec TTR 1 μM revealed ability to significantly increase the levels of IL-6, IL-8, Ang-2 and VEGF. Comparisons are relative to the negative control. (**B)** IL-6 levels measured by ELISA showed that while TTR 10 nM did not affect IL-6, TTR concentrations 25 nM-1 μM increased, in a concentration-dependent manner, the levels of IL-6. Data are expressed as mean ± SEM. * p<0.05; ** p<0.01; *** p<0.001.

We also confirmed the effect of IL-6 using an ELISA approach and as can be appreciated in Figure 3.B it is clear a dose-response effect as TTR concentration is increased. While at 10 nM the differences are not significant, TTR concentrations between 25 nM and 1μM lead to significantly increased expression of IL-6, as compared to the negative control. These results are also in line with those of the tube formation assay (Figure 2.A’).

### The impact of TTR reduction on BM thickening is greater in AD than in NT mice

To understand if TTR reduction impacts differently in an AD and in an non-AD environment, we analyzed the effect of the same TTR reduction on the collagen IV layer, in the AD and in the NT backgrounds (NT/TTR+/− versus AD/TTR+/−). Figure 4 depicts the results obtained and shows that the impact of TTR reduction is greater in an AD-like environment, adding relevance to TTR neuroprotection in AD. Given that these are 3-month-old animals and that, in this model, deposition begins at around 6 months, our results support other findings that suggest brain vascular dysregulation as the earliest factor during the disease progression.

**Figure 4.**
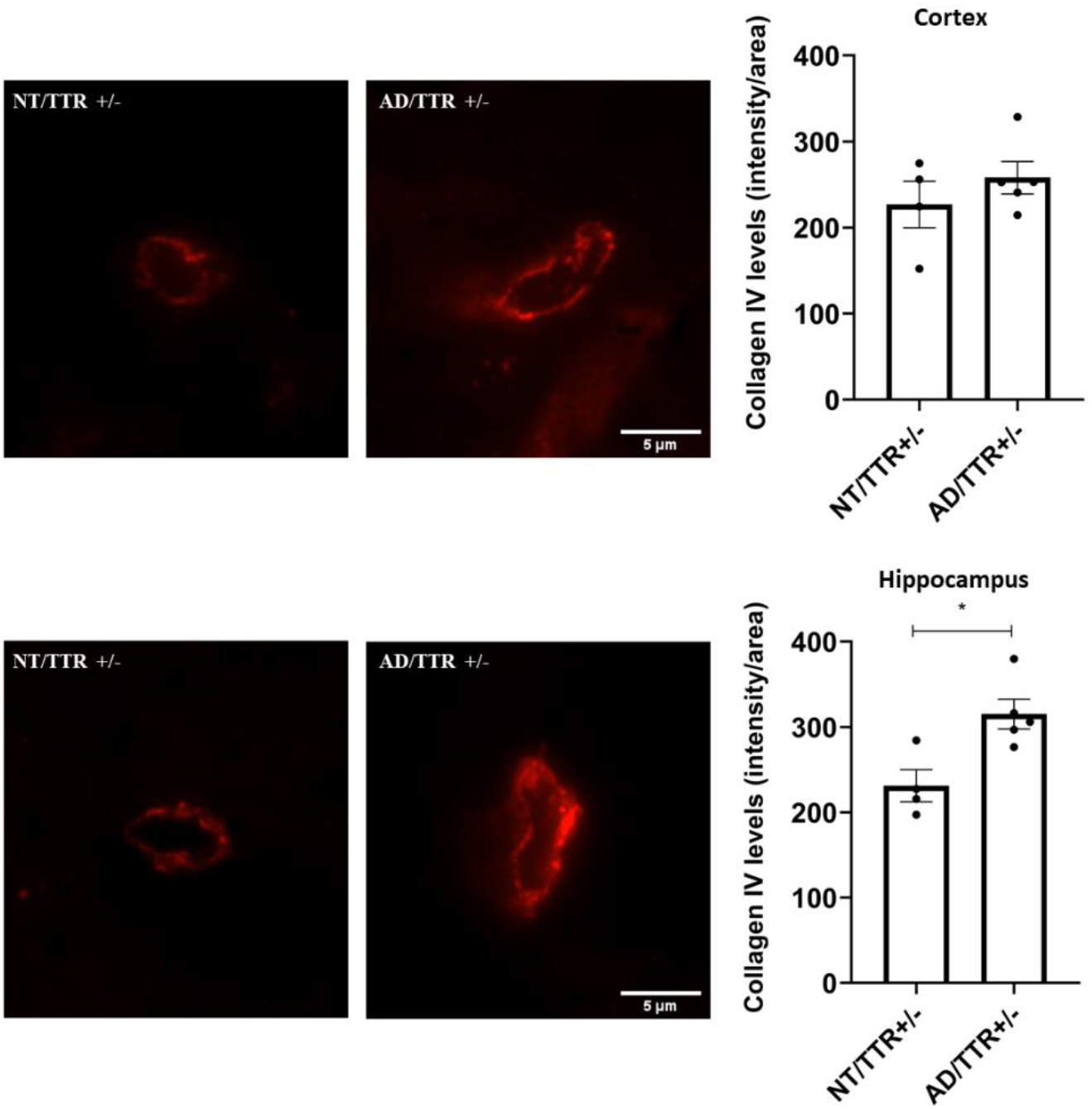
Effect of TTR reduction on collagen IV levels, in NT and AD mice. Representative images and quantification plots of collagen IV (red) immunostaining depicting microvessels of NT/TTR+/− (n=4) and AD/TTR+/− (n=5) 3-month-old mice in cortex and hippocampus, showing an increase in collagen IV expression in AD/TTR+/− mice relatively to NT/TTR+/− littermates, in the hippocampus. Data are expressed as mean ± SEM. * p<0.05. Scale bar = 5 μm.

### TTR stabilization results in decreased thickness of the collagen IV layer in brain microvessels of AD mice

So far, in this work, we have shown that TTR reduction worsens AD features in mice, as BM thickening. To further investigate a possible neuroprotective effect of TTR in the vascular context, and given that TTR stabilization is used to improve its activity, we analysed the BM thickness in brain microvessels in AD mice treated with one TTR stabilizer, IDIF.

Administration of IDIF to AD mice from the age of 5 to 7 month-old, resulted in amelioration of some AD features, such as the cognitive function and decreased Aβ brain levels [41].

In this work, we used brain slides obtained in our previous study, above mentioned [41] and performed collagen IV staining to assess BM thickness and vessels length in AD/TTR+/− animals, non-treated versus IDIF-treated. As depicted in Figure 5, AD/TTR+/− mice treated with IDIF presented a significant reduction in the BM thickness and increase in vessel length, as compared to non-treated animals. Altogether, these results indicate that TTR stabilization might be an avenue for early treatment in AD.

**Figure 5.**
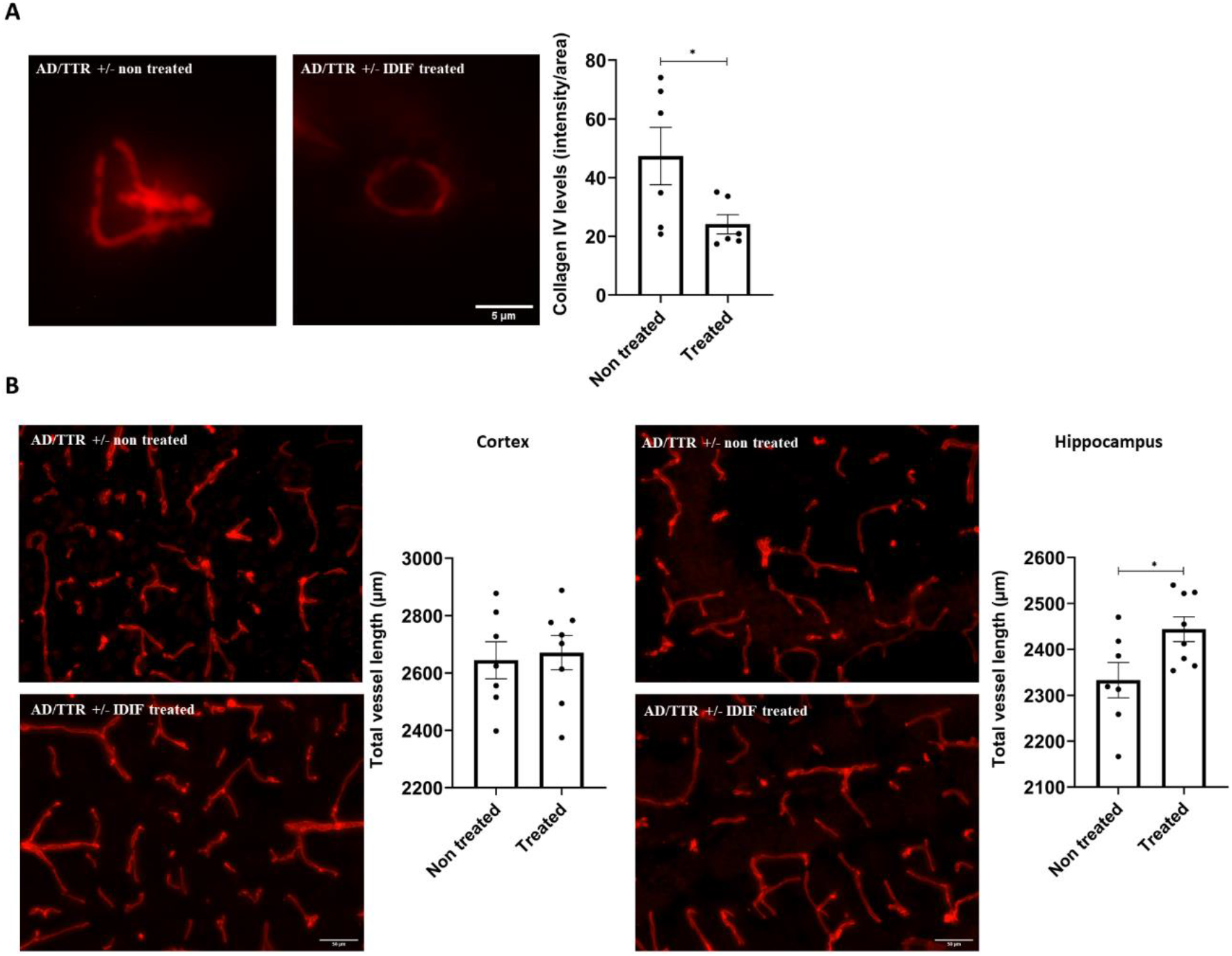
Effect of TTR stabilization by IDIF in the thickness of the collagen IV layer and vessel length of AD/TTR+/− mice. Representative images and quantification plots of vessels derived from AD/TTR+/− mice non-treated (n=6), or treated with IDIF (IDIF treated) (n=6) evidencing **(A)** a significantly decreased collagen IV layer in treated mice. Scale bar = 5 μm, and **(B)** a significantly increased vessel length in the hippocampus of IDIF treated compared to non-treated mice. Scale bar = 50 μm. Data are expressed as mean ± SEM. * p<0.05.

## Discussion

TTR is a homotetrameric protein typically known as a carrier of T4 and retinol in plasma and CSF. During the last years, several functions have been attributed to this protein, in particular, as a neuroprotector protein in physiologic and in disease contexts. In ischemia models, induced by permanent middle cerebral artery occlusion, TTR has been shown to be protective, as evidenced by the significant increase in cortical infarction, cerebral edema and the microglial-leukocyte response in mice with TTR deficiency compared with normal TTR levels [53]. Also, TTR deficiency results in spatial reference memory impairment [54]. Other works showed that TTR promotes nerve regeneration and axonal growth [55,56].

In AD, TTR binds to Aβ avoiding its aggregation, accumulation and toxicity, and facilitating its efflux across the BBB [26]. This barrier is essential to maintain brain homeostasis, however, during normal ageing and AD, BBB becomes dysfunctional contributing to disease progression. Molecules known to be important for Aβ brain homeostasis, such as LRP-1 and P-gp are reduced, and TTR was previously shown to increase LRP-1 expression in brain ECs and liver [26]. Thus, it is possible that TTR can regulate the neurovasculature in other ways, namely by influencing angiogenesis.

BM thickening through increased collagen IV levels is one such features observed in ageing and, more severely, in AD. Previous work by González-Marrero and co-workers described concomitant reduced TTR expression and thickening of the BM at the choroid plexus, in a triple transgenic mouse model of AD [57]. In addition, the authors reported increased Aβ42 in epithelial cytosol and in stroma surrounding choroidal capillaries [57]. Here, we showed that reduction of TTR expression in an AD mouse model influenced not only the BM, resulting in a thicker collagen IV layer, but also the vessel density, resulting in decreased vessel length. To ascertain if the differences were due to a direct or indirect effect of TTR, collagen IV levels were evaluated also in NT 3-month-old mice. NT/TTR+/− mice exhibited more collagen IV around brain microvessels than NT/TTR+/+ littermates, suggesting a direct effect of TTR.

In the AD model at the age of 3 months, Aβ deposition has not started. However, at the same age AD/TTR+/+ mice show decreased TTR compared to NT/TTR+/+ [49]. In addition, mild cognitive impaired patients show TTR decrease which further continues as AD progresses. Therefore, it is possible that TTR is early involved in the vascular alterations that occur in AD. Understanding the causes for TTR decrease and finding ways to restore its normal values and activity may be key in the development of a TTR-based therapy for AD.

It is not yet clear what leads to increased collagen IV levels in neurovasculature, but these changes are also found in rats suffering from chronic cerebral hypoperfusion [58,59], suggesting that decreased blood flow in the brain leads to high collagen IV content around the vessels. Indeed, diminished cerebral blood flow is an early impact event during AD development [1]. The thickened and rigid vascular wall may slow down nutrient supply and waste elimination, and possibly disturb perivascular drainage. This event along with the formed barrier will potentially contribute to progressive endothelial dysfunction and to an increasing accumulation of Aβ in the brain. We questioned if the effect of decreased TTR in the BM and vessel density could be related to TTR participation in angiogenesis, since a number of works implicated TTR in this process. These works suggest that TTR contributes to disease development by modulation ECs function. It is interesting to note that TTR levels are increased in those situations, such as diabetes type II, and in some reported situations of cancer, diseases where abnormal angiogenesis is an established hallmark [46,60,61]. TTR is decreased in AD [27–30] but its angiogenic potential was never evaluated *in vivo*. Using the *in vivo* CAM assay, we demonstrate that WT TTR, both produced recombinantly and purified from human plasma, influences angiogenesis by promoting the formation of new functional vessels.

As for the relation with brain angiogenesis, we showed that TTR promotes the formation of capillary-like structures by hCMEC/D3 cells and found that VEGF, Ang-2, IL-6 and IL-8 were significantly up-regulated in the presence of the protein. VEGF is a major driver of angiogenesis, playing a role in most of the steps of the process. Previous works have suggested a link between TTR and VEGF, and for example, elevated VEGF was found in the vitreous of patients with TTR amyloidosis [62]. Another work, proposed and interaction between the two molecules, although reporting that inhibition of VEGF in branch retinal vein occlusion (BRVO) upregulated TTR [63]. This can be interpreted as an attempt of the cells to restore VEGF levels by increasing TTR, thus corroborating our observations in hCMEC/D3 cells.

Ang-2 was previously found to be up-regulated in retinal ECs after treatment with TTR [44] and plays a controversial role in angiogenesis. If, on one hand, it increases migration capacity and tube formation in brain ECs [64], on the other hand, *in vivo* retinal studies showed that Ang-2 promotes EC death and vessel regression if VEGF is absent. However, when in the presence of VEGF it stimulates an increase in capillary diameter, remodeling of basal lamina, proliferation and migration of EC [65]. These studies support our findings where TTR promotes an increase of both VEGF and Ang-2, which should result in the promotion of angiogenesis. The observed upregulation of IL-8 and IL-6 is also consistent with an increase in angiogenesis given that IL-8 enhances proliferation, survival, migration and tube formation [66,67]; and IL-6 was shown to induce an increase in EC proliferation, migration and tube formation [68,69].

The importance of TTR in the AD pathogenesis is also patent when comparing AD with NT 3 months-old mice with the same TTR reduction, evidencing a thicker BM in the AD animals. Although this happens prior to amyloid-β deposition, we cannot exclude the presence of other Aβ species that can contribute to this increase, and add to the direct effect of TTR observed in the NT mice.

It is not known if the reduction of TTR in AD is a cause or effect of the disease but it is well known that TTR stability is key for its activity. Mutations in TTR, associated with amyloidosis, create tetrameric instability leading to dissociation into monomers. TTR stabilization seems also important to prevent pathological changes to the brain vasculature, and for example, heterozygous individuals with TTR T119M allele, which renders a more stable tetramer, have a reduced risk of cerebrovascular disease compared to homozygotes for WT TTR [70]. In AD, TTR stability is decreased, leading to accelerated clearance and consequently, to lower levels. We previously showed that TTR stabilization, achieved through the use of small-molecule compounds, sharing structural similarities with T4 and binding in the TTR central channel, results in improved TTR binding to Aβ [19]. One of those small-molecule stabilizers, IDIF administrated to our AD/TTR+/− mouse model resulted in the amelioration of AD features [40,41]. In this work we showed that IDIF reverted, at least partially, the vascular alterations induced by TTR decrease.

Our work provides positive and new results but it also reveals some limitations that should be mentioned. Concerning the studies with animals, only females were used, while a final conclusion on the effect of TTR decrease regarding the vascular alterations in AD may require the use of both genders. The animal model used in this study shows gender-associated modulation of brain Aβ levels by TTR, and females present a more severe AD-like neuropathology [49], which results in a more favourable scenario to assess the involvement of TTR in AD, explaining why we carried out our experiments in females.

Our results uncover angiogenesis as a mechanism in which TTR participates and importantly, it shows that TTR reduction has an impact in the vascular alterations that occur early in AD with the possibility of recovery upon TTR stabilization.

### Conclusions

In summary, this work shows that TTR has pro-angiogenic properties, up-regulating molecules such as IL-6, IL-8, Ang-2 and VEGF. TTR is also involved in the early vascular alterations occurring in AD, which may be used as a target for therapeutic intervention in AD.

## Abbreviations

AD: Alzheimer’s Disease
BM: basement membrane
TTR: Transthyretin
NT: non-transgenic
CAM: chick chorioallantoic membrane assay
IL: interleukin
Ang-2: angiopoietin-2
VEGF: vascular endothelial growth factor
IDIF: iododiflunisal
Aβ: amyloid-β peptide
LRP-1: low density lipoprotein receptor-related protein 1
BBB: blood-brain barrier
EC: endothelial cell
CSF: cerebrospinal fluid
FAP: Familial Amyloid Polyneuropathy
T4: thyroxine
bFGF: basic fibroblast growth factor
hCMEC/D3: human cerebral microvascular endothelial cell line
bEnd.3: mouse brain endothelial cell line.

## Declarations

### Funding

Grant from Norte2020, Portugal (Norte-01-0145FEDER-000008-Porto Neurosciences) and through a grant from Fundação Millennium bcp.

## Acknowledgments

The authors acknowledge the support of the i3S Scientific Platforms BioSciences Screening (BS) and Advanced Light Microcopy (ALM), members of the national infrastructure PPBI – Portuguese Platform of Bioimaging (PPBI-POCI-01-0145-FEDER-022122) and of the i3S Animal Facility. The flow cytometry analysis was performed at the Translational Cytometry i3S Scientific Platform with the assistance of E. Cardoso.

## Conflicts of Interest/Competing interests

The authors declare no conflict of interest.

## Ethics approval

All the above experiments were approved by the Institute for Research and Innovation in Health Sciences (i3s) Animal Ethics Committee and in agreement with the animal ethics regulation from Directive 2010/63/EU.

## Consent to participate

Not applicable

## Availability of data and material

All data and material present in this study available upon reasonable request to the corresponding author.

## Code availability

Not applicable

## Author Contributions

TG, JS and JRV performed the experiments; MP was responsible for the CAM assays, and respective analysis and data interpretation; IC was responsible for conception and supervision of the work. IC, TG, JS and GA discussed the results and wrote the manuscript. All authors reviewed the article and have read and agreed to the published version of the manuscript.

## Notes

### Competing Interest Statement

The authors have declared no competing interest.

## References

1. Iturria-Medina Y, Sotero RC, Toussaint PJ, Mateos-Pérez JM, Evans AC, Weiner MW, et al. Early role of vascular dysregulation on late-onset Alzheimer’s disease based on multifactorial data-driven analysis. Nat Commun. 2016;7:11934.

2. Donahue JE, Flaherty SL, Johanson CE, Duncan JA, Silverberg GD, Miller MC, et al. RAGE, LRP-1, and amyloid-beta protein in Alzheimer’s disease. Acta Neuropathol. 2006;112:405–15.

3. Vogelgesang S, Cascorbi I, Schroeder E, Pahnke J, Kroemer HK, Siegmund W, et al. Deposition of Alzheimer’s β-amyloid is inversely correlated with P-glycoprotein expression in the brains of elderly non-demented humans. Pharmacogenetics. 2002;12:535–41.

4. Daneman R, Prat A. The blood–brain barrier. Cold Spring Harb Perspect Biol. 2015;7.

5. Brown WR, Thore CR. Review: Cerebral microvascular pathology in ageing and neurodegeneration. Neuropathol Appl Neurobiol. 2011;37:56–74.

6. Kisler K, Nelson AR, Montagne A, Zlokovic B V. Cerebral blood flow regulation and neurovascular dysfunction in Alzheimer disease. Nat Rev Neurosci. 2017;18:419–34.

7. Biron KE, Dickstein DL, Gopaul R, Jefferies WA. Amyloid Triggers Extensive Cerebral Angiogenesis Causing Blood Brain Barrier Permeability and Hypervascularity in Alzheimer’s Disease. PLoS One. 2011;6.

8. de la Torre JC, Mussivand T. Can disturbed brain microcirculation cause Alzheimer’s disease? Neurol Res. England; 1993;15:146–53.

9. Paris D, Townsend K, Quadros A, Humphrey J, Sun J, Brem S, et al. Inhibition of angiogenesis by Aβ peptides. Angiogenesis. 2004;7:75–85.

10. Paris D, Patel N, DelleDonne A, Quadros A, Smeed R, Mullan M. Impaired angiogenesis in a transgenic mouse model of cerebral amyloidosis. Neurosci Lett. 2004;366:80–5.

11. Mancardi GL, Perdelli F, Rivano C, Leonardi A, Bugiani O. Thickening of the basement membrane of cortical capillaries in Alzheimer’s disease. Acta Neuropathol. 1980;49:79–83.

12. Merlini M, Meyer EP, Ulmann-Schuler A, Nitsch RM. Vascular β-amyloid and early astrocyte alterations impair cerebrovascular function and cerebral metabolism in transgenic arcAβ mice. Acta Neuropathol. 2011;122:293–311.

13. Thal DR, Capetillo-Zarate E, Larionov S, Staufenbiel M, Zurbruegg S, Beckmann N. Capillary cerebral amyloid angiopathy is associated with vessel occlusion and cerebral blood flow disturbances. Neurobiol Aging. 2009;30:1936–48.

14. Uspenskaia O, Liebetrau M, Herms J, Danek A, Hamann GF. Aging is associated with increased collagen type IV accumulation in the basal lamina of human cerebral microvessels. BMC Neurosci. 2004;5.

15. Bergström J, Gustavsson A, Hellman U, Sletten K, Murphy CL, Weiss DT, et al. Amyloid deposits in transthyretin-derived amyloidosis: cleaved transthyretin is associated with distinct amyloid morphology. J Pathol. England; 2005;206:224–32.

16. Schwarzman AL, Gregori L, Vitek MP, Lyubski S, Strittmatter WJ, Enghilde JJ, et al. Transthyretin sequesters amyloid beta protein and prevents amyloid formation. Proc Natl Acad Sci. 1994;91:8368–72.

17. Costa R, Gonçalves A, Saraiva MJ, Cardoso I. Transthyretin binding to A-Beta peptide – Impact on A-Beta fibrillogenesis and toxicity. FEBS Lett. 2008;582:936–42.

18. Du J, Murphy RM. Characterization of the interaction of β-Amyloid with Transthyretin monomers and tetramers. Biochemistry. 2010;49:8276–89.

19. Cotrina EY, Gimeno A, Llop J, Jiménez-Barbero J, Quintana J, Valencia G, et al. Calorimetric Studies of Binary and Ternary Molecular Interactions between Transthyretin, Aβ Peptides, and Small-Molecule Chaperones toward an Alternative Strategy for Alzheimer’s Disease Drug Discovery. J Med Chem. 2020;63:3205–14

20. Li X, Zhang X, Ladiwala ARA, Du D, Yadav JK, Tessier PM, et al. Mechanisms of transthyretin inhibition of β-amyloid aggregation in vitro. J Neurosci. 2013;33:19423–33.

21. Nilsson L, Pamrén A, Islam T, Brännström K, Golchin SA, Pettersson N, et al. Transthyretin Interferes with Aβ Amyloid Formation by Redirecting Oligomeric Nuclei into Non-Amyloid Aggregates. J Mol Biol. 2018;430:2722–33.

22. Palha JA, Moreira P, Wisniewski T, Frangione B, Saraiva MJ. Transthyretin gene in Alzheimer’s disease patients. Neurosci Lett. 1996;204:212–4.

23. Schwarzman AL, Gregori L, Vitek MP, Lyubski S, Strittmatter WJ, Enghilde JJ, et al. Transthyretin sequesters amyloid β protein and prevents amyloid formation. Proc Natl Acad Sci U S A. 1994;91:8368–72.

24. Schwarzman AL, Goldgaber D. Interaction of transthyretin with amyloid β-protein: Binding and inhibition of amyloid formation. CIBA Found Symp. 1996;146–64.

25. Cascella R, Conti S, Mannini B, Li X, Buxbaum JN, Tiribilli B, et al. Transthyretin suppresses the toxicity of oligomers formed by misfolded proteins in vitro. Biochim Biophys Acta – Mol Basis Dis. Elsevier B.V.; 2013;1832:2302–14.

26. Alemi M, Gaiteiro C, Ribeiro CA, Santos LM, Gomes JR, Oliveira SM, et al. Transthyretin participates in beta-amyloid transport from the brain to the liver – involvement of the low-density lipoprotein receptor-related protein 1? Sci Rep. Nature Publishing Group; 2016;6.

27. Ribeiro CA, Santana I, Oliveira C, Baldeiras I, Moreira J, Saraiva MJ, et al. Transthyretin Decrease in Plasma of MCI and AD Patients: Investigation of Mechanisms for Disease Modulation. Curr Alzheimer Res. 2012;9:881–9.

28. Han SH, Jung ES, Sohn JH, Hong HJ, Hong HS, Kim JW, et al. Human serum transthyretin levels correlate inversely with Alzheimer’s disease. J Alzheimer’s Dis. 2011;25:77–84.

29. Velayudhan L, Killick R, Hye A, Kinsey A, Güntert A, Lynham S, et al. Plasma transthyretin as a candidate marker for Alzheimer’s disease. J Alzheimer’s Dis. 2012;28:369–75.

30. Serot JM, Christmann D, Dubost T, Couturier M. Cerebrospinal fluid transthyretin: Aging and late onset Alzheimer’s disease. J Neurol Neurosurg Psychiatry. 1997;63:506–8.

31. Alemi M, Silva SC, Santana I, Cardoso I. Transthyretin stability is critical in assisting beta amyloid clearance– Relevance of transthyretin stabilization in Alzheimer’s disease. CNS Neurosci Ther. 2017;23:605–19.

32. Quintas A, Saraiva MJ, Brito RM. The amyloidogenic potential of transthyretin variants correlates with their tendency to aggregate in solution. FEBS Lett. England; 1997;418:297–300.

33. Almeida MR, Saraiva MJ. Clearance of extracellular misfolded proteins in systemic amyloidosis: Experience with transthyretin. FEBS Lett. 2012;586:2891–6.

34. Cardoso I, Goldsbury CS, Müller SA, Olivieri V, Wirtz S, Damas AM, et al. Transthyretin fibrillogenesis entails the assembly of monomers: a molecular model for in vitro assembled transthyretin amyloid-like fibrils. J Mol Biol. 2002;317:683–695.

35. Almeida MR, Gales L, Damas AM, Cardoso I, Saraiva MJ. Small transthyretin (TTR) ligands as possible therapeutic agents in TTR amyloidoses. Curr Drug Targets CNS Neurol Disord. Netherlands; 2005;4:587–96.

36. Bulawa CE, Connelly S, Devit M, Wang L, Weigel C, Fleming JA, et al. Tafamidis, a potent and selective transthyretin kinetic stabilizer that inhibits the amyloid cascade. Proc Natl Acad Sci U S A. 2012;109:9629–34.

37. Almeida MR, Macedo B, Cardoso I, Alves I, Valencia G, Arsequell G, et al. Selective binding to transthyretin and tetramer stabilization in serum from patients with familial amyloidotic polyneuropathy by an iodinated diflunisal derivative. Biochem J. 2004;381:351–6.

38. Baures PW, Oza VB, Peterson SA, Kelly JW. Synthesis and evaluation of inhibitors of transthyretin amyloid formation based on the non-steroidal anti-inflammatory drug, flufenamic acid. Bioorganic Med Chem. 1999;7:1339–47.

39. Miroy GJ, Lai Z, Lashuel HA, Peterson SA, Strang C, Kelly JW. Inhibiting transthyretin amyloid fibril formation via protein stabilization. Proc Natl Acad Sci U S A. 1996;93:15051–6.

40. Rejc L, Gómez-Vallejo V, Rios X, Cossio U, Baz Z, Mujica E, et al. Oral Treatment with Iododiflunisal Delays Hippocampal Amyloid-β Formation in a Transgenic Mouse Model of Alzheimer’s Disease: A Longitudinal in vivo Molecular Imaging Study. J Alzheimer’s Dis. Research Square; 2020;77:99–112.

41. Ribeiro CA, Oliveira SM, Guido LF, Magalhães A, Valencia G, Arsequell G, et al. Transthyretin stabilization by iododiflunisal promotes amyloid-β peptide clearance, decreases its deposition, and ameliorates cognitive deficits in an Alzheimer’s disease mouse model. J Alzheimer’s Dis. 2014;39:357–70.

42. Rios X, Gómez-Vallejo V, Martín A, Cossío U, Morcillo MÁ, Alemi M, et al. Radiochemical examination of transthyretin (TTR) brain penetration assisted by iododiflunisal, a TTR tetramer stabilizer and a new candidate drug for AD. Sci Rep. 2019;9:1–11.

43. Nunes RJ, De Oliveira P, Lages A, Becker JD, Marcelino P, Barroso E, et al. Transthyretin proteins regulate angiogenesis by conferring different molecular identities to endothelial cells. J Biol Chem. 2013;288:31752–60.

44. Shao J, Yao Y. Transthyretin represses neovascularization in diabetic retinopathy. Mol Vis. 2016;22:1188–97.

45. Shao J, Yin Y, Yin X, Ji L, Xin Y, Zou J, et al. Transthyretin Exerts Pro-Apoptotic Effects in Human Retinal Microvascular Endothelial Cells Through a GRP78-Dependent Pathway in Diabetic Retinopathy. Cell Physiol Biochem. 2017;43:788–800.

46. Lee C-C, Ding X, Zhao T, Wu L, Perkins S, Du H, et al. Transthyretin Stimulates Tumor Growth through Regulation of Tumor, Immune, and Endothelial Cells. J Immunol. 2019;202:991–1002.

47. Borchelt DR, Ratovitski T, Van Lare J, Lee MK, Gonzales V, Jenkins NA, et al. Accelerated Amyloid Deposition in the Brains of Transgenic Mice Coexpressing Mutant Presenilin 1 and Amyloid Precursor Proteins. Neuron. 1997;19:939–45.

48. Episkopou V, Maeda S, Nishiguchi S, Shimada K, Gaitanaris GA, Gottesman ME, et al. Disruption of the transthyretin gene results in mice with depressed levels of plasma retinol and thyroid hormone. Proc Natl Acad Sci U S A. 1993;90:2375–9.

49. Oliveira SM, Ribeiro CA, Cardoso I, Saraiva MJ. Gender-dependent transthyretin modulation of brain amyloid-β Levels: Evidence from a mouse model of Alzheimer’s disease. J Alzheimer’s Dis. 2011;27:429–39.

50. Furuya H, Saraiva MJM, Alves IL, Gawinowicz MA, Saraiva MJM, Alves IL, et al. Production of Recombinant Human Transthyretin with Biological Activities toward the Understanding of the Molecular Basis of Familial Amyloidotic Polyneuropathy (FAP). Biochemistry. 1991;30:2415–21.

51. Almeida MR, Damas AM, Lans MC, Brouwer A, Saraiva MJ. Thyroxine binding to transthyretin Met 119: Comparative studies of different heterozygotic carriers and structural analysis. Endocrine. 1997;6:309–15.

52. Santos LM, Rodrigues D, Alemi M, Silva SC, Ribeiro CA, Cardoso I. Resveratrol administration increases transthyretin protein levels, ameliorating AD features: The importance of transthyretin tetrameric stability. Mol Med. 2016;22:597–607.

53. Santos SD, Lambertsen KL, Clausen BH, Akinc A, Alvarez R, Finsen B, et al. CSF transthyretin neuroprotection in a mouse model of brain ischemia. J Neurochem. 2010;115:1434–44.

54. Sousa JC, Marques F, Dias-Ferreira E, Cerqueira JJ, Sousa N, Palha JA. Transthyretin influences spatial reference memory. Neurobiol Learn Mem. 2007;88:381–5.

55. Fleming CE, Saraiva MJ, Sousa MM. Transthyretin enhances nerve regeneration. J Neurochem. 2007;103:831–9.

56. Fleming CE, Mar FM, Franquinho F, Saraiva MJ, Sousa MM. Transthyretin internalization by sensory neurons is megalin mediated and necessary for its neuritogenic activity. J Neurosci. 2009;29:3220–32.

57. González-Marrero I, Giménez-Llort L, Johanson CE, Carmona-Calero EM, Castañeyra-Ruiz L, Brito-Armas JM, et al. Choroid plexus dysfunction impairs beta-amyloid clearance in a triple transgenic mouse model of Alzheimer’s disease. Front Cell Neurosci. 2015;9:1–10.

58. Ueno M, Tomimoto H, Akiguchi I, Wakita H, Sakamoto H. Blood-brain barrier disruption in white matter lesions in a rat model of chronic cerebral hypoperfusion. J Cereb Blood Flow Metab. United States; 2002;22:97–104.

59. De Jong GI, Farkas E, Stienstra CM, Plass JRM, Keijser JN, De La Torre JC, et al. Cerebral hypoperfusion yields capillary damage in the hippocampal CA1 area that correlates with spatial memory impairment. Neuroscience. 1999;91:203–10.

60. Zemany L, Bhanot S, Peroni OD, Murray SF, Moraes-Vieira PM, Castoldi A, et al. Transthyretin Antisense Oligonucleotides Lower Circulating RBP4 Levels and Improve Insulin Sensitivity in Obese Mice. Diabetes. United States; 2015;64:1603–14.

61. Mody N, Graham TE, Tsuji Y, Yang Q, Kahn BB. Decreased clearance of serum retinol-binding protein and elevated levels of transthyretin in insulin-resistant ob/ob mice. Am J Physiol – Endocrinol Metab. 2008;294:785–93.

62. O’Hearn TM, Fawzi A, He S, Rao NA, Lim JI. Early onset vitreous amyloidosis in familial amyloidotic polyneuropathy with a transthyretin Glu54Gly mutation is associated with elevated vitreous VEGF. Br J Ophthalmol. 2007;91:1607–9.

63. Cehofski LJ, Kruse A, Alsing AN, Nielsen JE, Pedersen S, Kirkeby S, et al. Intravitreal bevacizumab upregulates transthyretin in experimental branch retinal vein occlusion. Mol Vis. Molecular Vision; 2018;24:759–66.

64. Mochizuki Y, Nakamura T, Kanetake H, Kanda S. Angiopoietin 2 stimulates migration and tube-like structure formation of murine brain capillary endothelial cells through c-Fes and c-Fyn. J Cell Sci. 2002;115:175–83.

65. Lobov IB, Brooks PC, Lang RA. Angiopoietin-2 displays VEGF-dependent modulation of capillary structure and endothelial cell survival in vivo. Proc Natl Acad Sci U S A. 2002;99:11205–10.

66. Dwyer J, Hebda JK, Le Guelte A, Galan-Moya EM, Smith SS, Azzi S, et al. Glioblastoma Cell-Secreted Interleukin-8 Induces Brain Endothelial Cell Permeability via CXCR2. PLoS One. 2012;7.

67. Li A, Dubey S, Varney ML, Dave BJ, Singh RK. IL-8 Directly Enhanced Endothelial Cell Survival, Proliferation, and Matrix Metalloproteinases Production and Regulated Angiogenesis. J Immunol. 2003;170:3369–76.

68. Fee D, Grzybicki D, Dobbs M, Ihyer S, Clotfelter J, MacVilay S, et al. Interleukin 6 promotes vasculogenesis of murine brain microvessel endothelial cells. Cytokine. 2000;12:655–65.

69. Hernández-Rodríguez J, Segarra M, Vilardell C, Sánchez M, García-Martínez A, Esteban MJ, et al. Elevated production of interleukin-6 is associated with a lower incidence of disease-related ischemic events in patients with giant-cell arteritis: Angiogenic activity of interleukin-6 as a potential protective mechanism. Circulation. 2003;107:2428–34.

70. Hornstrup LS, Frikke-Schmidt R, Nordestgaard BG, Tybjcrg-Hansen A. Genetic stabilization of transthyretin, cerebrovascular disease, and life expectancy. Arterioscler Thromb Vasc Biol. 2013;33:1441–7.

